# Brain-Derived CCN3 Is An Osteoanabolic Hormone That Sustains Bone in Lactating Females

**DOI:** 10.1101/2023.08.28.554707

**Authors:** Muriel E. Babey, William C. Krause, Candice B. Herber, Kun Chen, Zsofia Torok, Joni Nikkanen, Ruben Rodriquez, Xiao Zhang, Fernanda Castro-Navarro, Yuting Wang, Saul Villeda, Nancy E. Lane, Erica L. Scheller, Charles K.F. Chan, Thomas H. Ambrosi, Holly A. Ingraham

## Abstract

In lactating mothers, the high calcium (Ca^2+^) demand for milk production triggers significant bone resorption. While estrogen would normally counteract excessive bone loss and maintain sufficient bone formation during this postpartum period, this sex steroid drops precipitously after giving birth. Here, we report that brain-derived CCN3 (Cellular Communication Network factor 3) secreted from KISS1 neurons of the arcuate nucleus (ARC^KISS1^) fills this void and functions as a potent osteoanabolic factor to promote bone mass in lactating females. Using parabiosis and bone transplant methods, we first established that a humoral factor accounts for the female-specific, high bone mass previously observed by our group after deleting estrogen receptor alpha (ERα) from ARC^KISS1^ neurons^1^. This exceptional bone phenotype in mutant females can be traced back to skeletal stem cells (SSCs), as reflected by their increased frequency and osteochondrogenic potential. Based on multiple assays, CCN3 emerged as the most promising secreted pro-osteogenic factor from ARC^KISS1^ neurons, acting on mouse and human SSCs at low subnanomolar concentrations independent of age or sex. That brain-derived CCN3 promotes bone formation was further confirmed by in vivo gain- and loss-of-function studies. Notably, a transient rise in CCN3 appears in ARC^KISS1^ neurons in estrogen-depleted lactating females coincident with increased bone remodeling and high calcium demand. Our findings establish CCN3 as a potentially new therapeutic osteoanabolic hormone that defines a novel female-specific brain-bone axis for ensuring mammalian species survival.

## MAIN TEXT

Osteoporosis significantly impacts healthy aging and is commonly experienced by more women than men. Females leverage estradiol (E2) to increase energy expenditure^2^ and preserve bone mass^3^ by regulating bone remodeling via osteocytes^4^, osteoblasts^5^, and osteochondral skeletal stem cells (ocSSCs)^6^, which are fated for bone and cartilage^7,8^. For women, natural or drug-induced estrogen depletion following menopause or anti-hormone therapies slowly degrades bone mass, underscoring the anabolic features of estrogen on bone metabolism. However, the intimate association between estrogen and bone is oddly uncoupled during lactation when the late-stage pregnancy E2 surge drops precipitously. Bone remodeling also rises sharply to supply the high calcium demand by progeny (reviewed in ^9^). Parathyroid hormone-related protein (PTHrP), a close orthologue of parathyroid hormone (PTH) from mammary glands, is the main driver for stripping calcium from maternal bones for milk production^10,11^. The relentless demand for calcium by newborns eventually leads to significant bone loss in mothers, dropping nearly 30% in rodents due to large litter sizes^12^ and 10% in humans^13,14^; these losses mostly normalize post-lactation^12,14^. Presumably, the maternal skeleton (and that of pups) would be severely compromised without this concomitant lactational-anabolic phase, as inferred by the increased bone mass in lactating mothers after conditional knock-out of PTHrP^15,16^. Absent the underlying mechanism driving the anabolic arm of increased bone remodeling during lactation, this notion remains to be proved.

Aside from the direct actions of E2 on bone, we and others have shown that central estrogen signaling exerts a sex-dependent restraint on bone formation, alongside its role in promoting spontaneous activity and thermogenesis^17-20^. High bone mass in females results following the deletion of neuronal ERa^21^ in the arcuate nucleus of the medial basal hypothalamus (MBH)^1 22^. Viral and genetic mouse models designed to eliminate ERα in the ARC, specifically in ARC^Kiss1^ neurons, resulted in high trabecular bone mass in the spine and long bones, independent of high E2 levels, confirming the central origins of this extraordinary bone phenotype^1^.

Here, using a combination of question-driven and discovery-based approaches, we set out to identify the osteogenic hormone-like factor responsible for the high bone mass in mutant females after first showing that this factor circulates in the blood. Cellular communication network factor 3 (CCN3/Nov) emerged as the best candidate, fulfilling all anticipated criteria; it is secreted, its appearance in the ARC coincides with the onset and then loss of the bone phenotype after a dietary challenge, it causes new bone formation when ectopically delivered ex vivo to bones or in vivo to mice, and finally, bone mass degrades in mutant females after knockdown of Ccn3 transcripts in the ARC. We propose that CCN3 functions as a neuroendocrine hormone in a newly-identified brain-bone axis evolved to sustain skeletal health in mammalian mothers and offspring.

### A Humoral Factor Mediates High Bone Mass in Mutant Females

Our prior genetic and viral deletions of ERα in the ARC strongly suggested that a subset of KNDy neurons in the ARC regulates bone mass and bone strength in females but not males (Fig 1a,b)^1^. That KNDy neurons participate in this brain-bone axis was further supported after deleting ERα with the Prodynorphin-Cre driver (Fig. 1b and Extended Data Fig. 1a,b). To identify the molecular origins of the high bone mass phenotype, we relied exclusively on the *Es-r1*^*Nkx2-1Cre*^ female mouse model, which exhibits this unusual phenotype by four weeks of age (Extended Data Fig. 1c-e). Given the privileged position of the ARC as one of the few circumventricular organs of the brain just dorsal to the median eminence, we asked if this high bone mass in mutant females might originate from a circulatory factor.

**Figure 1.**
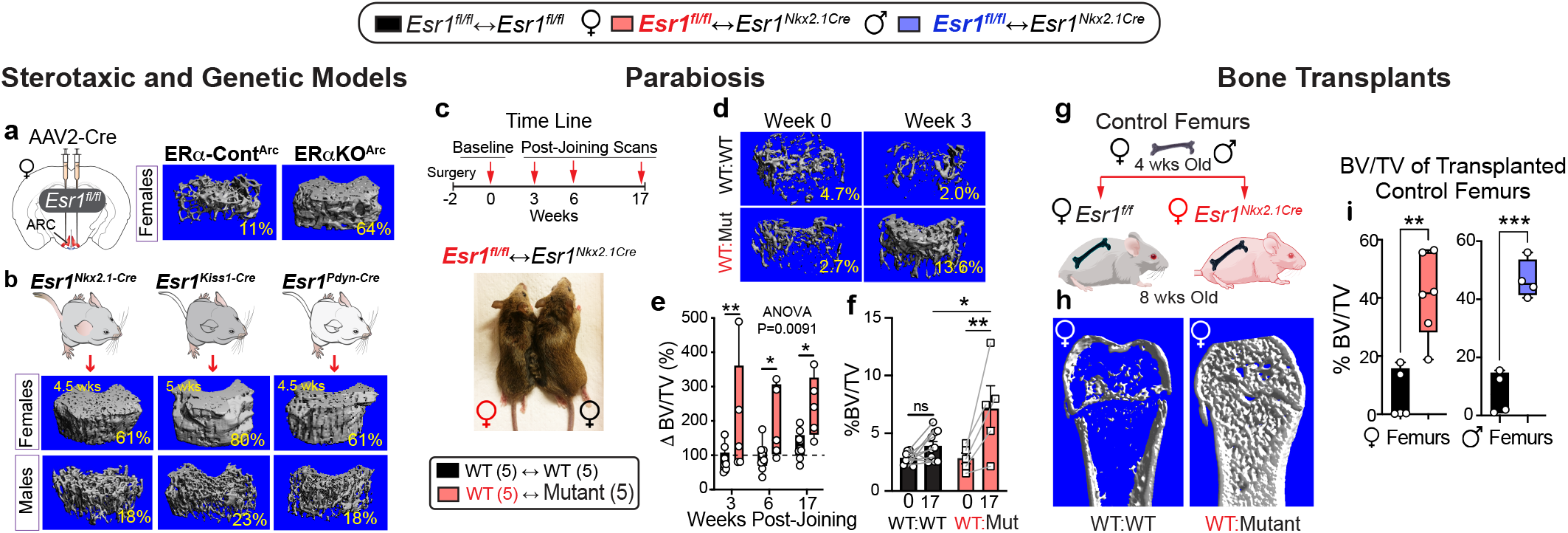
A Brain-Dependent Circulatory Factor Functions as An Osteoanabolic Hormone. **a**, Schematic showing stereotaxic deletion of ERa in the ARC using AAV2-Cre vector with representative μCT scans of the distal femur from females injected with AAV2 control virus (ERa-Control^ARC^ female) and ERa-KO^ARC^ female, as previously reported^1^. **b**, μCT images obtained from *Esr1*^*Nkx2*.*1-Cre*^, *Esr1*^*Kiss-Cre*,^ and *Esr1*^*Pdyn-Cre*^ 4.5-5-week-old mice with %BV/TV indicated lower righthand corner. **c**, Timeline of in vivo μCT imaging post-surgical pairing of *Esr1*^*fl/fl*^ (WT) and *Esr1*^*Nkx2*.*1-Cre*^ (Mutant) female mice. **d**, Representative in vivo μCT imaging of distal femur at baseline (Wk 0) and 3 weeks later (Wk 3) with %BV/TV indicated. **e**, Bar graph shows the percent change in %BV/TV at Wks 3, 6, and 17 compared to Wk 0. **f**, Absolute %BV/TV graphed for *Esr1*^*fl/fl*^ females in the WT:WT (black) and WT:Mutant pairs (red) showing values for each animal at baseline (0) and 17 weeks later (17), (N = 5WT:5WT, 5WT:5Mut). **g**, Schematic of wildtype female and male bone transplants into *Esr1*^*fl/fl*^ (WT) and *Esr1*^*Nkx2*.*1-Cre*^ (Mutant) female mice for 6 weeks. **h**, Representative image of μCT scan for control femur transplanted into WT (WT:WT) or mutant females (WT:Mutant). **i**, Fractional bone volume of excised bones transplanted into *Esr1*^*fl/fl*^ females (black bars) or female (red bar) or male (blue bar) bones transplanted into *Esr1*^*Nkx2*.*1-Cre*^ females (N = 4-6). Two-way ANOVA in left panel e with repeated measures, and One-way ANOVA in right panel f (Šidák’s multiple-comparisons test). Unpaired Student’s T-test, 2-tailed in panel i. *p < 0.05, **p < 0.01, ***p < 0.001, ns = not significant. Error Bars ± SEM.

Using classical parabiosis coupled with in vivo μCT imaging (Fig. 1c), two female groups were surgically joined, generating control pairs (WT-WT) and controls joined to mutant females (WT-MUT). Shortly after surgery (2 wks), baseline bone structural parameters on the contralateral femur opposite the surgical side were established for each animal in the pairings. As predicted, females in the WT-WT paring exhibited a net decrease in bone mass that was readily observed beginning at six weeks post-surgery (Fig. 1e); this decline normalizes by Wk 17, rising an average of ∼37%. In the WT-MUT pairings, higher fractional bone volume (%BV/TV) was observed at all time points in control females, increasing ∼152% by 17 Wks (Fig. 1f and Extended Data Fig. 2a,b). Uterine weights, as well as other parameters, were unchanged between WT-WT and WT-MUT pairings consistent with the notion that higher estrogen levels are not at play in generating high bone mass in mutant females (Extended Data Fig. 2c). We noted that the already high bone mass in mutant females increased further in some WT-MUT pairings (Extended Data Fig. 2d,e).

Bone transplant studies confirmed that a humoral factor accounts for bone mass in mutant females. Female and male femurs from 4-week-old control donors were implanted subcutaneously into 8-week-old wild-type or mutant females (Fig. 1g and Extended Data Fig. 3a). Significant increases in fractional bone mass were detected in both female and male control femurs six weeks post-implantation as determined by μCT analyses (Fig. 1h,i and Extended Data Fig. 3b,c), arguing strongly that this brain-dependent, female-specific osteoanabolic hormone will function in both sexes.

### A Brain-Dependent Factor Alters Skeletal Stem Cell Dynamics and Bone Anabolic Activity

Skeletal homeostasis is tightly regulated by skeletal stem cell (SSC)-based bone formation and osteoclast-based bone resorption. Chan, Ambrosi, and colleagues demonstrated that stem cells with distinct lineage hierarchies and cellular contributions facilitate new bone formation^23^. In particular, osteochondral SSCs (ocSSCs) form bone and cartilage (and fibro/stromal lineage cell populations but not bone marrow adipocytes) and are present in the growth plate and periosteum of bones^8,24^ (Fig. 2a), whereas perivascular SSCs (pvSSCs) give rise to unilateral committed adipogenic progenitor cells (APCs) that generate all bone marrow adipose tissue (BMAT) in bones^23,25^. As such, we reasoned that the brain-dependent anabolic hormone might alter ocSSC activity, given the increased bone formation in mutant females^1^.

**Figure 2.**
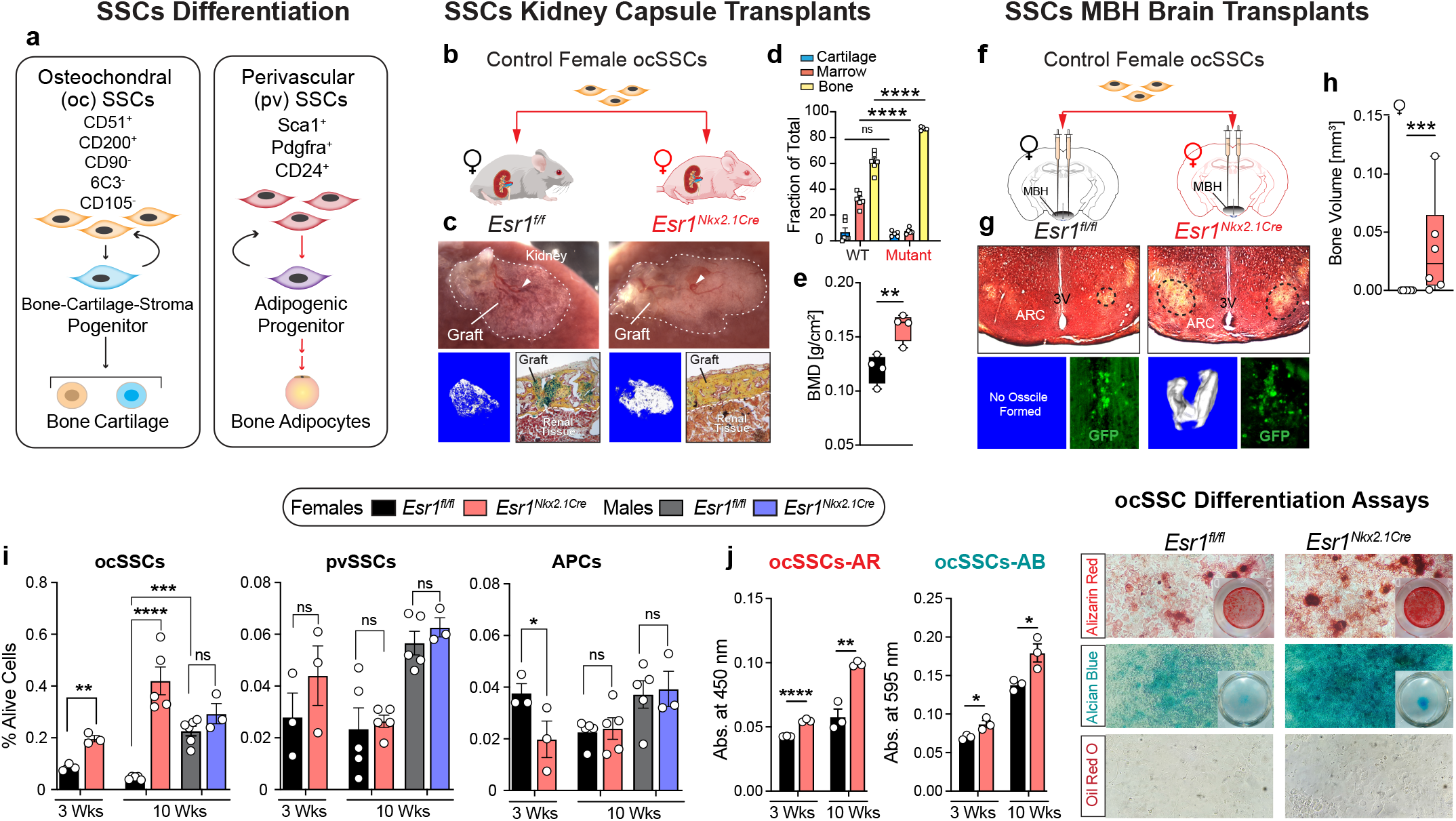
Brain-Dependent Bone Factor Increases Osteogenic Capacity of ocSSCs. **a**, Schematic of FACS-isolation and the fate of ocSSCs (left panel) and pvSSCs (right panel) using listed cell-surface markers. **b**, Schematic of wild-type female ocSSCs (∼15,000 live cells) transplanted into kidney capsule of *Esr1*^*fl/fl*^ and *Esr1*^*Nkx2*.*1-Cre*^ female mice. **c**, Images of outlined graft region with host-derived hematopoietic features (top panels, white arrows). Volumetric bone density images (left panels) and sections of graph region stained for mineralized bone (yellow), cartilage (blue), and marrow (red) in right panels). **d**, Fractional areas for marrow, cartilage, and bone quantified from stained sections in individual kidney grafts (N = 6, 5), and **e**, bone density for a separate cohort (N = 4, 4). **f**, Schematic of stereotaxic bilateral delivery of FACS-purified control ocSSCs (∼550 live cells) from *Esr1*^*fl/fl-CAG-Luc,-GFP*^ into the MBH of *Esr1*^*fl/fl*^ and *Esr1*^*Nkx2*.*1-Cre*^ female mice. **g**, Representative images of Pentachrome (top panel) with ossicles from imaged brains (lower left) and anti-GFP (lower right) of stained brain sections, six weeks post-injection. **h**. Measured volumetric bone density of ossicles in MBH of *Esr1*^*fl/fl*^ (black) and *Esr1*^*Nkx2*.*1-Cre*^ (red) female mice. **i**, Percent of FACS-purified ocSSCs, pvSSCs, APCs described in Methods isolated from *Esr1*^*fl/fl*^ and *Esr1*^*Nkx2*.*1-Cre*^ 3 weeks old (N = 3, 3) and 10-week-old females (N = 5, 5) and males (5, 3) with the legend for bar graphs above. **j**, Isolated ocSSCs pooled from 3- and 10-week-old females differentiated in defined media and stained with Alizarin Red (left panel) or Alcian Blue (right panel) with brightfield images from representative wells including staining with Oil Red O (n = 3, 3 per group). One-way ANOVA in panels d, i, and j (Šidák’s multiple-comparisons test). Unpaired Student’s T-test, 2-tailed for panels e, h, and i (3 Wks). *p < 0.05, **p < 0.01, ***p < 0.001, ****p < 0.0001, ns = not significant. Error Bars ± SEM.

OcSSCs from female wild-type mice were isolated by flow cytometry and transplanted beneath the renal capsule of control or mutant females (Fig. 2b). As expected, wild-type ocSSCs transplanted into control *Esr1*^*fl/fl*^ female littermates formed an ectopic bone graft with a host-derived hematopoietic compartment over six weeks (Fig. 2c,d). However, wild-type ocSSCs grafted into *Esr1*^*Nkx2-1Cre*^ females exhibited significantly higher mineralization and little hematopoietic marrow (Fig. 2c-e), suggesting that the brain-dependent osteoanabolic hormone present in mutant females alters the ocSSC lineage to promote bone formation (Fig. 2e). Consistent with this hypothesis, the higher fractional bone mass in WT-MUT parabionts or whole bone transplants studies correlated with increased ocSSCs frequency (Extended Data Fig. 4a). The robustness of this putative circulatory osteoanabolic hormone was further verified by stereotaxic delivery of GFP-positive wild-type ocSSCs to the vicinity of the ARC (Fig. 2f). Remarkably, μCT imaging of mutant *Esr1*^*Nkx2-1Cre*^ hypothalami revealed mineralized ossicles overlapping with transplanted GFP+ cells six weeks post-injection; no ossicles were detected in wild-type brains (Fig. 2g,h). These data further establish the existence of a circulatory anabolic bone factor in mutant females, perhaps originating from the ARC or surrounding hypothalamic regions.

That a circulating factor might promote wild-type ocSSC activity prompted us to compare the differentiation capacity of mutant and control ocSSCs. Flow cytometric analysis revealed a sex-dependent increased frequency of ocSSCs in both pre-pubertal (3Wks) and young adult mutant (10Wks) females (Fig. 2i). This alteration was limited to ocSSCs, as pvSSCs and their progeny adipogenic progenitor cells (APCs) fated for BMAT^23,25^ were equivalent in controls and mutants, except for the lower frequency of APCs in younger mutants (Fig. 2i). Differentiation assays revealed that while ocSSCs from both genotypes showed no differences in colony-forming ability (CFU-F) (Extended Data Fig. 4b), mutant ocSSCs exhibit a high intrinsic potential for bone and cartilage formation (Fig. 2j). Enhanced ocSSCs activity was also observed in older *Esr1*^*Nkx2-1Cre*^ females, consistent with their attenuated bone loss compared to age-matched control littermates and the known linkage between ocSSC dysfunction and age-related bone loss (Extended Data Fig. 4c-e)^26^.

### Identification of CCN3 as a Candidate Brain-Derived Osteogenic Factor

Despite the expansion and enhanced osteogenic capacity of mutant ocSSCs, single-cell RNA sequencing (sc-RNAseq) data implied that while mutant ocSSCs differentiation dynamics were primed towards bone formation, only modest overall transcriptomic differences were detected; thus, unfortunately, providing few hints regarding the identity of the osteoanabolic hormone in *Esr1*^*Nkx2-1Cre*^ females (Extended Data Fig. 5). However, the first helpful clue in our hunt for this anabolic bone hormone arose after finding that a chronic high-fat dietary (HFD) challenge dramatically disrupts the sex-dependent brain-bone axis. While body weights, fat mass, blood triglycerides, and glucose homeostasis remained unchanged with HFD (Extended Data Fig. 6a), the high bone mass phenotype in *Esr1*^*Nkx2-1Cre*^ females completely reversed with this dietary challenge (Fig. 3a). Reduction in trabecular and cortical bone mass was accompanied by expected structural changes and secondary spongiosa (Extended Data Fig. 6b) as well as reduced bone strength (Fig 3a,b). Histomorphometry showed that osteoclast number, bone formation, and mineralized surface remained proportional to bone surface in mutant females fed HFD (Fig. 3b and Extended Data Fig. 6c,d). We noted that levels of BMAT, quantified by osmium staining^27^, were significantly lower in mutant females on either SD or HFD compared to control littermates (Fig 3a, b). Thus, while dense bones in *Esr1*^*Nkx2-1Cre*^ females readily degrade with HFD, they fail to accumulate BMAT to a similar extent as control females, thus defying the normal linkage between BMAT expansion and bone loss^28^. Bone parameters in *Esr1*^*Nkx2-1Cre*^ males remained unchanged from their control littermates (Extended Data Fig. 6e,f). Moreover, chronic hyperglycemia induced by the insulin receptor antagonist S961^29^, failed to degrade mutant female bone mass, demonstrating the specificity of HFD-induced bone loss in mutant females (Extended Data Fig. 6g).

**Figure 3.**
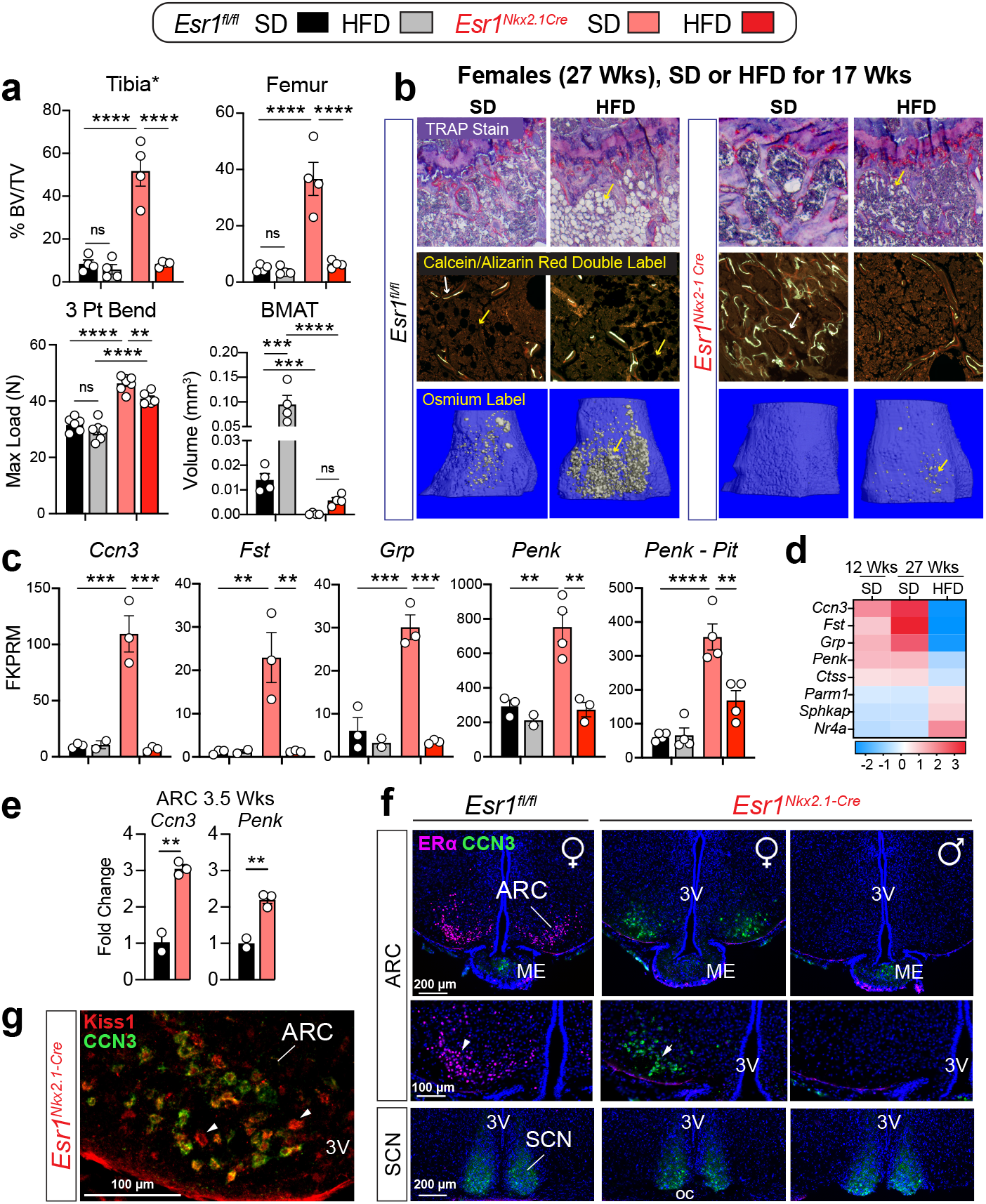
Identification of CCN3 as a Candidate Brain-Derived Osteoanabolic Factor. **a**, Fractional bone volume, mechanical strength, and BMAT levels in long bones of Esr1^fl/fl^ and Esr1^Nkx2.1-Cre^ females fed SD or HFD for 17 weeks (N = 4-6 per group). **b**, Representative images of tibia stained for TRAP (top row), labeled for Calcein/ Alizarin Red (middle row) and osmium stained (bottom row); double labeling (white arrows), lipid droplets (yellow arrows). **c**, Normalized reads for candidate genes; Penk in the pituitary (N = 2-4). **d**, Heatmap of top DEGs changed in Esr1^N-kx2.1-Cre^ females at 12 weeks of age (adapted from RE), and at 27 weeks of age fed SD or HFD; Scale is Log2 fold. **e**, Transcript levels of Ccn3 and Penk in mutant female ARC by qPCR. **f**, Staining for ERa (pink) and CCN3 (green) in brain sections from posterior ARC and SCN regions of Esr1^fl/fl^ female (10 wks) and Esr1^Nkx2.1-Cre^ female and male (12 wks), Scale bar = 100 and 200 μm. **g**, Merge of CCN3 and KISS1 expression in Esr1^Nkx2.1-Cre^ female ARC. One-way ANOVA in panels a and c (Šidák’s multiple-comparisons test). Un-paired Student’s T-test, 2-tailed for panels e. **p < 0.01, ***p < 0.001, ****p < 0.0001, ns = not significant. Error Bars ± SEM. Abbreviations: ME median eminence, ARC arcuate nucleus, 3V third ventricle, SCN suprachiasmatic nucleus, oc optic chiasm.

We then exploited the dynamic changes in bone following HFD by profiling gene changes in microdissected ARC from *Esr1*^*Nkx2-1Cre*^ females fed SD or HFD. Bulk RNA-seq revealed a small set of upregulated differentially expressed genes (DEGs) encoding neuropeptides or secreted proteins in the ARC, including *Ccn3, Fst, Grp*, and *Penk* – all dropped significantly after HFD but were elevated on SD in young and older females (Fig. 3c,d and Extended Data Fig. 7a). Importantly, *Ccn3* and *Penk* expression correlated well with the onset of the high bone mass at around four weeks of age (Fig. 3e) but were unchanged at earlier time points or in *Esr1*^*Nkx2-1Cre*^ males (Extended Data Fig. 7b,c). Except for Penk in the mutant pituitary, few notable candidates emerged after profiling the pituitary and liver, two common sources of secreted proteins, (Fig. 3c and Extended Data Fig. 7a). CCN3/ *Ccn3* expression was nearly absent in control females or males but readily detected in the basal region of the mutant female ARC colocalizing with KISS1, a known marker of KNDy neurons (Fig. 3f,g); expression levels easily surpassed those detected in the suprachiasmatic nucleus (SCN)^30^.

### CCN3 Functions as an Osteoanabolic Bone Hormone

High expression of CCN3 in mutant ARC neurons that are KISS1-positive and ERα-negative, prompted us to test this founding member of the CCN family^31^. This secreted protein is postulated to antagonize CCN2 to inhibit osteogenesis^32,33^, although a single report suggests the opposite^34^. Nevertheless, the anabolic potential of CCN3 was evaluated in three distinct assays. First, ex vivo whole long bone cultures were treated with CCN3 after initially finding that plasma collected from *Esr1*^*Nkx2-1Cre*^ females elevated bone mass of freshly dissected control femurs after five days, rising an average of 37% and 50% for female and male femurs, respectively (Fig. 4a,b and Extended Data Fig. 8a). Importantly, long bones of both sexes showed substantial degradation from their baseline values when cultured in media alone (Extended Data Fig. 8b,c). Using this simple but effective assay, we found that low doses of mouse (m)CCN3 (0.31 nM) induced an upward dynamic shift (60%) in bone mass compared to saline (Fig. 4d-f and Extended Data Fig. 8b). Second, adult wild type mice were injected with mCCN3 (i.p.) or saline daily over three weeks. Within this short time and at this low dose (7.5μg/kg), a significant increase in bone mass was noted for both female and male mice treated with mCCN3 (Fig. 4g,h and Extended Data Fig. 8d). Finally, culturing primary ocSSC isolated from newborn wild-type mice treated daily with mCCN3 increased mineralization by ∼200% (Fig. 4i). Met-ENK and BAMP-22, two major peptides encoded by Penk, failed to elicit any changes; these negative results are consistent with the fact that the competitive antagonist of the μ-opioid receptor, Naloxone, was unable to degrade the high bone mass in mutant female mice (Extended Data Fig. 9a). Additionally, low concentrations of human CCN3 increased osteogenesis in primary human ocSSCs from pubertal and aged/geriatric patients independent of sex; higher levels had no beneficial effect on osteogenesis and were inhibitory at the highest level (Fig. 4j and Extended Data Fig. 9b-e). The pro-osteogenic effect on young and old human ocSSCs appears to be exclusive to CCN3 (Extended Data Fig. 9b, c). Taken together, these data suggest that at low doses, CCN3 is an anabolic bone hormone in mice and humans.

**Figure 4.**
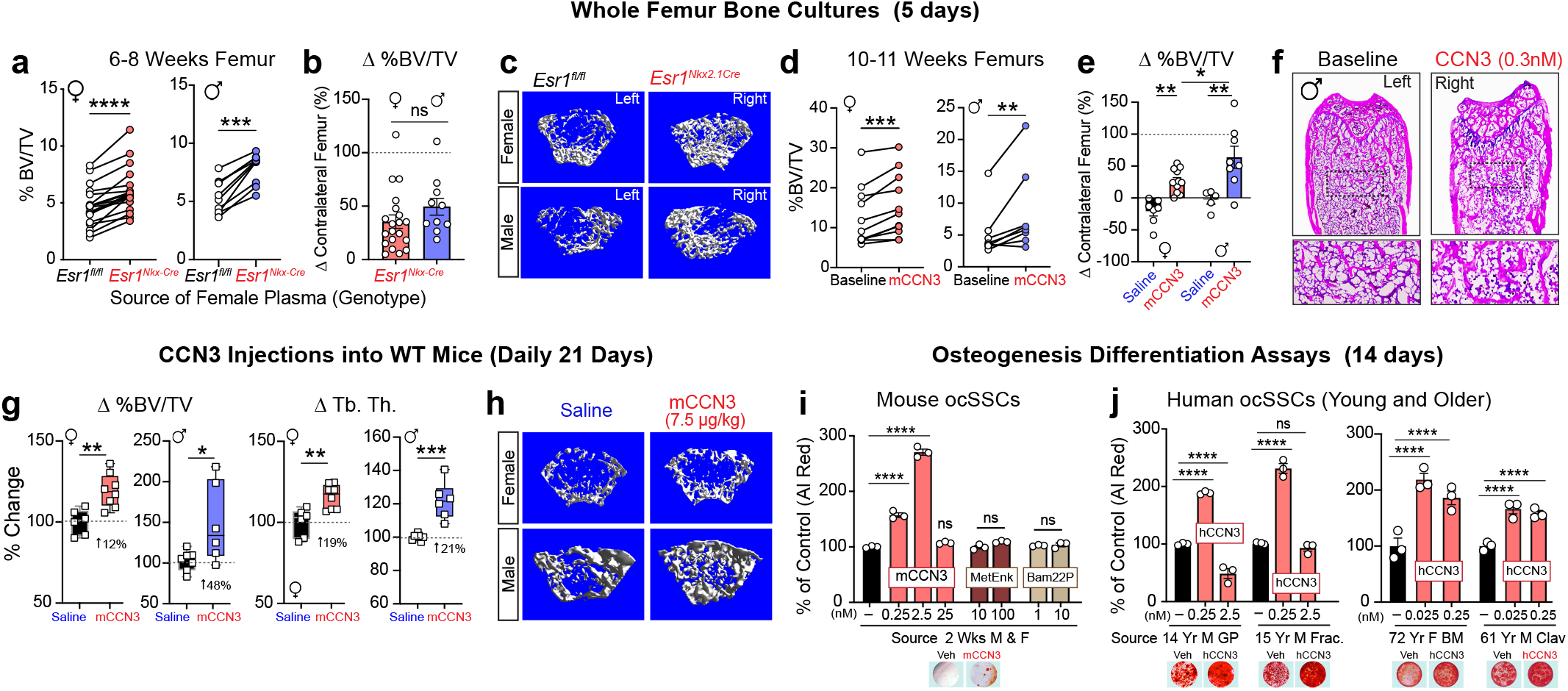
CCN3 Independent of Sex- and Age Promotes New Bone Formation Through Enhanced Osteogenesis of ocSSCs in Mice and Humans. **a**, Fractional bone volume for control Esr1^fl/fl^ female (red circles) and male (blue circles) femurs cultured ex vivo and treated daily for 5 days with plasma isolated from Esr1^Nkx2.1-Cre^ mutant females (6-12 wks); contralateral femurs were treated with control plasma (white circles), (N = 19, 10). **b**, Data from panel a, showing percent change in bone volume of contralateral female (red bar) or male (blue bar) femurs after adding mutant versus control plasma. **c**, Representative μCT images scanned from female and male femurs treated with plasma with corresponding %BV/TV. **d**, Fractional bone volume of female (N =11) and male (N=8) control femurs treated daily with recombinant mouse CCN3 (mCCN3, 0.3 nM) compared to untreated baseline control (Baseline). **e**, Percent change in bone volume of female (red bar) or male (blue bar) femurs treated with either CCN3 or saline and then compared to the baseline value of freshly isolated and stored contralateral femur (N =5-10). **f**, Representative images of H&E-stained femurs at baseline and after culturing with CCN3. **g**, Percent change in %BV/TV and trabecular thickness in Esr1^fl/fl^ females and males following daily CCN3 injections (i.p. 7.5 μg/kg) or saline for 21 days. All data were normalized to the average for control females injected with only saline (N = 6,8 for females and 7,6 for males). **h**, Representative μCT images scanned from female and male treated femurs with corresponding %BV/TV. **i**, Osteogenic differentiation assays of purified mouse ocSSCs treated with recombinant mouse (m)CCN3 protein or the Penk-encoded met-ENK and Bam22P peptides. Plotted values are normalized to Alizarin staining of control wells in defined media alone, set at 100% with brightfield images of wells below without or with mCCN3 (n = 3 per condition). **j**, Osteogenic differentiation assays of purified human ocSSCs treated with recombinant human (h)CCN3 protein. Plotted values are normalized to Alizarin staining of control wells in defined media alone set at 100%, and representative images of wells treated without or with mCCN3 below (n = 3 per condition). One-way ANOVA in panels e, k, and i (Šidák’s multiple-comparisons test). Paired Student’s T-test, 2-tailed for panels a, d and Unpaired Student’s T-test for panels b, g. *p < 0.05, **p < 0.01, ***p < 0.001, ****p < 0.0001, ns = not significant. Error Bars ± SEM.

### Brain-Derived CCN3 Is Used During Lactation to Preserve Bone Mass

To establish the linkage between CCN3 and increased bone density in *Esr1*^*Nkx2-1Cre*^ females, bone parameters were measured following overexpression or knockdown of CCN3 in control or mutant females, respectively. We leveraged the secretory capacity of hepatocytes to increase circulating CCN3 in *Esr1*^*fl/fl*^ control females after systemic delivery of the AAV-dj-CCN3 vector with high liver tropism^35^. Ectopic CCN3 expression in liver hepatocytes was detected as early as two weeks post-injection (Fig. 5a and Extended Data Fig. 10a). Even at exceedingly low levels of hepatic CCN3 expression, modest increases in bone formation rate were observed (Extended Data Fig 10a,b). At 10-fold higher levels of hepatic CCN3 expression, fractional bone volume was enhanced by nearly 80% (Fig. 5b,c and Extended Data Fig. 10b). Conversely, transient knockdown of Ccn3 expression in *Esr1*^*N-kx2-1Cre*^ female ARC neurons by stereotaxic delivery of siRNAs attenuated the dense bone phenotype with levels of Ccn3/ CCN3 tracking well with %BV/TV (Fig. 5d-f). Together, these data show that elevating or lowering CCN3 levels in control and mutant females, respectively, affects bone mass and that maintenance of the high bone mass in *Esr1*^*Nkx2-1Cre*^ females relies on sufficient CCN3 in ARC^ERα-CCN3^ neurons.

**Figure 5.**
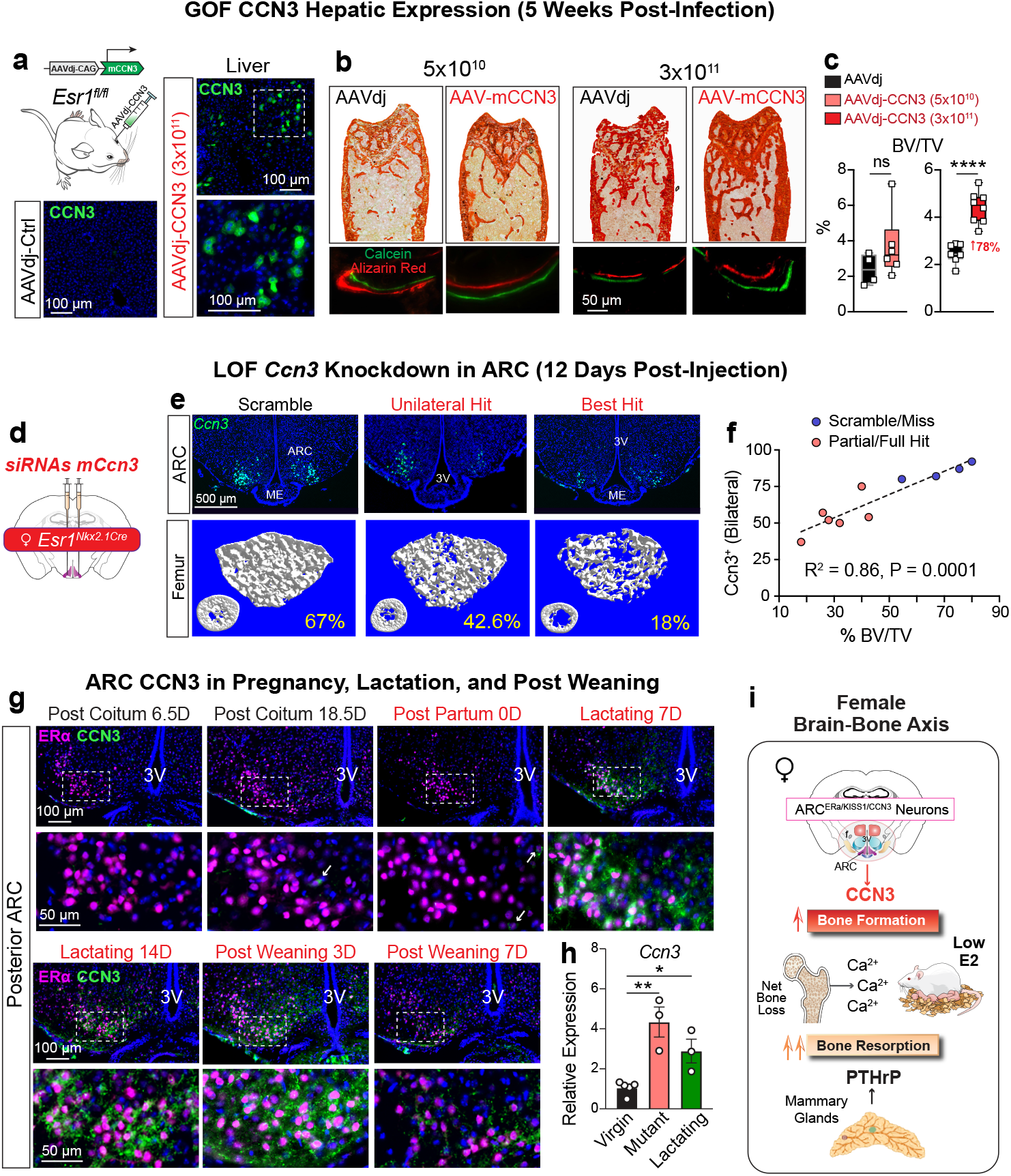
Brain-Derived CCN3 Drives Higher Bone Mass and Is Activated in Lactating Females When Estrogen a Plummets. **a**, Ectopic expression of mCCN3 in hepatocytes with their characteristic double nuclei following retroorbital injection of AAVdj-CAG-CCN3 compared to injections of control AAVdj viral vector (AAVdj-Ctrl), Scale bar = 100 μm. **b**, Representative images of femur sections stained with Alizarin Red and magnified area of double-labeled bone surfaces obtained from control females injected with 100 μL of low (5*10^10^ GC/mL) or high (3*10^11^ GC/mL) AAVdj-CAG-CCN3 dose of viral vector. **c**, BV/TV (%) from μCT and bone formation rate (BFR) plotted as determined by dynamic histomorphometry analyses and assessed by Calcein (green) and Alizarin Red (red). Low dose (N = 4,6, High Dose N = 7,8). **d**, Injection of siRNA directed against *mCcn3* into ARC of mutant *Esr1*^*Nkx2*.*1-Cre*^ mutant females. **e**, Ccn3 expression in posterior ARC shown for scramble control oligos, a unilateral hit, and the best level of knockdown. Scale bar = 100 μm. Lower panels show corresponding μCT scans of sagittal and longitudinal views of distal femur; complete misses were added to the control group (N = 4, 4). **f**, XY plot of the number of CCN3-postivie neurons in the posterior ARC versus the fractional bone volume. **g**, Representative images of coronal female brain sections stained for ERα (pink) and CCN3 (green) in the posterior ARC region during pregnancy or different postpartum stages as indicated; higher resolution images are shown in the lower row (N ≥ 2). Scale bar = 100 μm upper row, 50 μm lower row. **h**, *Ccn3* transcripts quantified from microdissected ARC tissue obtained from from *Esr1*^*fl/fl*^ virgin, *Esr1*^*Nkx2*.*1-Cre*^ virgin, and *Esr1*^*fl/fl*^ lactating (7DPP) females (N= 5, 3, 3). **I**, Schematic of brain-derived CCN3 as an osteoanabolic hormone that counteracts the catabolic actions of mammary-gland PTHrP to maintain adequate calcium and preserve the maternal skeleton during lactation when E2 levels drop. One-Way ANOVA for panel g (Šidák’s multiple-comparisons test), Unpaired Student’s T-test for panels b. *p < 0.05, **p < 0.01, ****p < 0.0001, ns = not significant. Error Bars ± SEM. Abbreviations: ME median eminence, ARC arcuate nucleus, 3V third ventricle.

To expand our findings beyond an artificial genetic phenomenon, we asked if CCN3 appears at any life stage in the ARC of control females, focusing on the postpartum period when maternal BFR escalates to maintain the skeletal calcium reservoir^12,36^, in the face of exceedingly low circulating E2^9^. Similar to virgin controls, CCN3-expressing ARCERa neurons are non-existent in early and late-stage pregnancy (Fig. 5g,h). However, by seven days postpartum (DPP),CCN3 is abundantly expressed in a subset of ARC^ERα^ neurons of lactating dams (Fig. 5g), reaching near equivalent levels as found in *Esr1*^*Nkx2-1Cre*^ mutant females (Fig. 5h). Forced weaning reduced CCN3 in ARC^ERα^ neurons when examined 3 or 7 days after removal of pups (10 or14 DPP, respectively), suggesting that the need for bone-promoting CCN3 lessens at the cessation of lactation when calcium demand diminishes (Fig. 5g). These data imply that brain-derived CCN3 in ARC^ERα-KISS1-CCN3^ neurons sustains sufficient bone formation to support inter-generational resource transfer when ovarian-derived estrogens fade away (Fig 5i).

## DISCUSSION

The role of ARC^ERα-KISS1^ neurons as the gatekeeper of female reproduction is well-established – these neurons control multiple facets of physiology, including regulating pubertal onset and the hypothalamic-pituitary-gonadal axis. Here, we discover yet another crucial function for ARC^ERα-KISS1^ neurons in females – controlling bone homeostasis during lactation via the brain-derived osteoanabolic hormone CCN3. Shutting down ovarian estradiol production and energy expenditure during lactation^37^poses a serious problem - how do mineralized bone surfaces keep up while being “plundered” for calcium during lactation^9^, especially in the trabecular-rich spine, which is particularly susceptible to lactational-associated osteoporosis^13^ as well as the loss of ERa signaling in bone cells^4^. Through CCN3, ARC^ERα-KISS1-CCN3^ neurons solve this problem by lifting the usual restraints on energetically costly bone formation, thereby increasing trabecular bone in the spine and long bones1, which degrades during lactation. How does CCN3 become upregulated in the female ARC during this unique life stage? While Ccn3 could be coordinately upregulated along with Kiss1 due to the sharp rise in prolactin signaling^38^, we note that prolactin levels are only modestly elevated in mutant females1. In addition to bone homeostasis, it will be of interest to determine if the medial basal hypothalamus also directly or indirectly controls other adaptive responses, including the dramatic increases in intestinal length and nutrient absorption in lactating mothers^39^.

Our data are at odds with the proposed function of CCN3 as an inhibitor of osteogenesis, as reviewed in^40^. However, most prior studies relied on high doses and high overexpression of CCN3, making it likely that compensatory cellular responses or non-specific receptor activation in cellular bone niches account for the adverse effects on osteogenesis. We, too, observed inhibitory effects in both mouse and human ocSSC differentiation assays at CCN3 doses exceeding the sub-nanomolar range. Based on the presence of anti-parallel b-strands and the C-terminal Cystine knot (CTCK) domain that mediates disulfide-linked dimerization^41^, we predict that CCN3 circulates at low doses as a tightly held homodimer, binding its cognate receptor with high affinity, similar to other growth factors such as NGF. Identifying the molecular target of CCN3 in ocSSCs and possibly other cellular populations, including osteocytes that reversibly remodel their perilacunar/canalicular matrix during lactation^42^, will help resolve these discrepancies. We speculate that permanently deleting ERα from ARC^KISS1^ neurons leads to continuous secretion of low levels of CCN3, recapitulating the anabolic phase of healthy, postpartum bone remodeling. CCN3, in combination with E2 and absent PTHrP, rapidly generates strong, dense bones. Whether our findings can be translated into new therapeutics for mitigating osteoporosis and fracture risk remains an exciting future direction.

## METHODS

### Ethics

Experiments were approved and performed in accordance with the guidelines of the UCSF Institutional Animal Care Committee (IACUC) or the UCD Animal Ethics Committee, the National Institutes of Health Guide for Care and Use of Laboratory Animals, and recommendations of the International Association for the Study of Pain. The ages of animals used in this study ranged between one to seventy weeks old and included both male and female mice.

### Mice

The origin of the *Esr1*^*fl/fl*^ allele (official allele: *Esr1tm1Sakh*) on a 129P2 background and used to generate *Esr1*^*Nkx2-1Cre*^ mice are described1 and were maintained on CD-1;129P2 mixed background. Primer sequences used for genotyping are listed in Extended Data Table 1. *Esr1*^*Nkx2-1Cre-CAG-Luc,-GFP*^ mice were generated by crossing male mice harboring the *CAG-Luc-GFP* allele (official allele: *L2G-85Chco/J*) to female mice homozygous for the *Esr1*^*fl/fl*^ allele, followed by an additional cross to generate *Esr1*^*fl/fl*^; ^*Nkx2-1Cre*^; ^*Luc-GFP*^ colony, which was maintained on a mixed FVB/N, CD-1, 129P2, and C57BL/6 genetic background. *Esr1*^*ProdynorphinCre*^ mice were generated by crossing homozygous *Esr1fl/fl* females to *Prodynorphin-Cre* (B6;129S-*Pdyntm1*.*1(cre)Mjkr*/LowlJ, purchased from JAX) males. Unless otherwise noted, animals were given ad-lib access to a standard lab diet (Pico Lab, 5058, LabDiet, 4kcal% fat) and sterile water and housed under controlled and monitored rooms for temperature and humidity with a 12h light/dark cycle. All animal procedures were performed in accordance with UCSF institutional guidelines under the Ingraham lab IACUC protocol of record.

### Parabiosis

Parabiosis surgery followed previously described procedures^43^. Briefly, 6-week-old *Esr1*^*fl/fl*^ and *Esr1*^*Nkx2-1Cre*^ females underwent mirror-image incisions at the left and right flanks through the skin, and incisions were made through the abdominal wall. The peritoneal openings of the adjacent parabionts were sutured together. Additionally, elbow and knee joints from each parabiont were sutured together. The skin incision of each mouse was then stapled. Each mouse was injected subcutaneously with enrofloxacin (Baytril, Bayrer) antibiotic and buprenorphine (Butler Schein) and monitored during surgical recovery. To monitor the health of pairs, their body weights and grooming behaviors were monitored weekly. The pairing was checked by the presence of Evans Blue dye in blood collected 2 hrs post-injection of 200 μL of 0.5% Evans Blue from the submandibular of the uninjected parabiont.

In vivo micro (μ) CT was performed to determine changes in trabecular bone mass over time using a Scanco Viva CT40 high-speed μCT preclinical scanner. Prior to scanning, mice were anesthetized and placed in a 3D-printed nose cone to accommodate surgically paired mice. Distal femurs of each mouse in the parabiont group (opposite to the incision side) were imaged 2 weeks post-surgery at baseline (0) and then at 3, 6, and 17 weeks from baseline imaging at intervals that preserve bone mass^44^. Parabionts were anesthetized, and a 2 mm region of the contralateral distal femur (opposite to the surgically paired side) for assessing the trabecular bone compartment of 1 mm length proximal to the epiphyseal plate and cortical parameters at the diaphysis in an adjacent 0.4 mm region of the femur. Imaging was performed by the UCSF Skeletal Biology and Biomechanics Core supported by the NIAMS Award P30AR075055.

### Bone and ocSSC Kidney and Brain Transplants

Briefly, long bones were dissected from 4-week-old female or male *Esr1*^*fl/fl*^ mice, cleaned of excess tissue, and immediately implanted under the skin after creating a horizontal incision and small pocket posterior to the scapula in 8-week-old acceptor *Esr1*^*fl/fl*^ or *Esr1*^*Nkx2-1Cre*^ female mice. After 6 weeks of incubation, donor femurs were subsequently removed and analyzed by ex-vivo μCT. Briefly, volumetric bone density and volume were measured at the distal femur using a Scanco Medical μCT 50 specimen scanner calibrated to a hydroxyapatite phantom or Bruker SkyScan1276 (Bruker Preclinical Imaging). For Sanco imaging, samples were fixed in 10% phosphate-buffered formalin and scanned in 70% ethanol using a voxel size of 10 mm and an X-ray tube potential of 55 kVp and X-ray intensity of 109 μA. Scanned regions included a 2 mm region of the femur proximal to the epiphyseal plate. For Bruker Skyscan 1276 imaging, a source voltage of 85 kV, a source current of 200 μA, a filter setting of AI 1 mm, and a pixel size of 12-20 microns (a set number was used for samples of a specific experiment) at 2016 × 1344 were used. Reconstructed samples were analyzed using CT Analyser and CTvox software (Bruker). Standard best practices were used to quantify trabecular and cortical bone parameters^44^. Image acquisition of an implanted femur was captured with iPhone 13 Pro and then edited in Photoshop CC.

For kidney transplant assays, primary ocSSCs were isolated as described below from 4-week-old *Esr1*^*fl/fl*^ female mice. Approximately 20,000 live cells were resuspended in 2 μL Matrigel (Brand), and the entire mixture was injected into the renal capsule of 8-week-old recipient mice. Six weeks post-transplantation, animals were then euthanized, and kidney grafts were processed as described below. Excised kidneys with renal capsule grafts and brain tissue with injected stem cell grafts were scanned using the same settings. For the processing of kidneys after SSC transplants, kidneys were dissected out and cleaned of soft tissue, fixed in 4% PFA, and embedded in OCT for cryosectioning. Sections (5 μm) were subsequently stained with standard Movat’s Pentachrome staining kit (Abcam Cat# ab245884). Brightfield images were taken using a Luminera Infinity-3 and quantified using ImageJ software. Excised kidneys containing renal capsule grafts injected with SSCs were scanned using the same settings as above for bone transplant assays.

For brain transplants, *Esr1*^*fl/fl-CAG-Luc,-GFP*^ ocSSCs were isolated by FACS as described below and kept on ice in sterile artificial cerebral spinal fluid (Tocris Bioscience, Bristol, UK cat#3525). Approximately 2 hours after cell isolation, ∼400-700 ocSSCs in 1 μL of solution were delivered bilaterally above the 3rd ventricle by the ARC nucleus by stereotaxic injection (A/P: -1.58; M/L: +-0.3; D/V: -5.95 from Skull) into 12–18-week-old *Esr1*^*fl/fl*^ or *Esr1*^*Nkx2-1Cre*^ females. Six weeks post-implant, mice were perfused, and immunohistochemistry was performed on cryosections (20 μm) collected from brains fixed in 4% paraformaldehyde using standard procedures. GFP staining used a polyclonal chicken anti-GFP antibody (Novus Biologicals, Littleton, CO cat# NB100-1614) at 1:2500. Images were taken using the Keyence B2-X800. Excised medial basal hypothalamic brain tissues from female mice injected with stem cells were scanned using the same settings as described above for bone transplant assays.

### Flow Cytometry Isolation of Primary Skeletal Stem Cells

Flow cytometry and cell sorting were performed on a FACS Aria II cell sorter (BD Biosciences) and analyzed using FlowJo software. Mouse long bones and human patient fracture callus samples (IRB35711; SCRO-525) were dissected and freed from the surrounding soft tissue, which was then followed by dissociation with mechanical and enzymatic steps as described^45,46^. Briefly, the tissue was placed in collagenase digestion buffer supplemented with DNase and incubated at 37°C for 60 min under constant agitation. After collagenase digestion and neutralization, undigested materials were gently triturated by repeated pipetting. Total dissociated cells were filtered through a 70-m nylon mesh and pelleted at 200 *×g* at 4°C for 5 min. Cells were resuspended in ACK (ammonium-chloride-potassium) lysing buffer to eliminate red blood cells and centrifuged at 200 *×g* at 4°C for 5 min. The pellet was resuspended in 100 μl staining media (2% FBS/phosphate-buffered saline [PBS]) and stained with antibodies for at least 30 min at 4°C (antibody information can be found in Table X). Living cells were gated for lack of PI (propidium iodide; 1:1000 diluted stock solution: 1 μg/ml in water; mouse cells) signal or DAPI (human cells). Compensation, fluorescence-minus-one control-based gating, and FACS isolation were conducted prior to analysis or sorting using established antibody cocktail combinations. A complete list of antibodies used for FACS-purification of SSCs can be found in Extended Data Table 2.

For mouse SSC lineages, antibodies used were as follows: CD90.1 (Thermo Fisher, cat. no. 47–0900-82), CD90.2 (Thermo Fisher, Cat#47–0902-82), CD105 (Thermo Fisher, Cat#13–1051-85), CD51 (BD Biosciences, Cat#551187), CD200 (BD Biosciences, Cat#745255), CD45 (BioLegend, Cat#103110), Ter119 (Thermo Fisher, Cat#15–5921-81), Tie2 (Thermo Fisher, Cat#14–5987-81) 6C3 (BioLegend, Cat#108312), and Streptavidin PE-Cy7 (Thermo Fisher, Cat#25–4317-82) as well as Sca-1 (Thermo Fisher, Cat#25–5981), CD45 (Thermo Fisher, Cat#11–0451), CD31 (Thermo Fisher, Cat#11–0311), CD140a (ThermoFisher, Cat#17–1401), CD24 (Thermo Fisher, Cat#47–024).

For human SSC isolation, antibodies used were as follows: CD45 (BioLegend, Cat#304029-BL), CD235a (BioLegend, Cat#306612-BL), CD31 (Thermo Fisher Scientific, Cat#13-0319-82), CD202b (TIE-2) (BioLegend, Cat#334204), CD146 (BioLegend, Cat#342010), PDPN (Thermo Fisher Scientific, Cat#17-9381-42), CD164 (BioLegend, Cat#324808), and CD73 (BioLegend, Cat#344016).

### Cell Culturing and Differentiation Assays of Primary Mouse and Human SSCs

Only freshly sorted primary murine or human ocSSCs were used in this study. After cell isolation by FACS, primary cells were cultured as described above. Mouse cells were cultured in minimum essential medium alpha (MEM-α) with 10% FBS and 1% penicillin-streptomycin (ThermoFisher; Cat#15140–122) and maintained in an incubator at 37°C with 5% CO2. Human cells were cultured in MEM-alpha medium (Fisher Scientific: Cat#12561-056) with 10% human platelet-derived lysate (Stem Cell Technologies, Cat#06960) and 1% Penicillin-Streptomycin solution (Pen-Strep; Thermo Fisher Scientific, Cat#15140-122). To induce osteogenic differentiation, pre-confluent cells were supplemented with osteogenesis-inducing factors 100 nM dexamethasone, 0.2 mM L-ascorbic acid 2-phosphate, and 10 mM β-glycerophosphate for 14 days.

For testing of candidate factors [mCcn3 (NovusBiological, Cat#NBP2-35100), hCCN3 (NovusBiologicals, Cat#NBP2-35084), mFst (NovusBiological, Cat# NBP2762685U), hFST (Stemcell Technologies, Cat#50-197-6487), Bam22P (Sigma, Cat# SCP0057), MetEnk (Sigma, Cat#M6638), hGRP (RayBiotech, Cat#230-00695-10], indicated concentrations were added to defined media and changed every 2nd day with fresh media. Cells were then formalin-fixed and stained with 2% Alizarin red S (Roth) in distilled water. Wells were washed twice with PBS and once with distilled water. Oil red O staining was performed by fixing cells with 4% PFA for 15 min at room temperature using an oil red O working solution prepared from a 0.5% stock solution in isopropanol and was diluted with distilled water at a ratio of 3:2. The working solution was filtered and applied to fixed cells for at least 1 hour at room temperature. Cells were washed four times with tap water before evaluation. CFU-F assays were conducted by freshly sorting a defined number of cells of desired cell populations into separate culture dishes containing expansion media. The medium was changed twice a week. cells were fixed and stained with crystal violet (Sigma) on day 10 of culturing.

### HFD Challenge and Metabolic and Bone Parameters

High-fat diet was purchased from Research Diets (D12492, 60 kcal% Fat). Glucose tolerance tests were conducted after a 6-hour fast with glucose administered intraperitoneally (1.0 g/kg of body mass). Blood glucose and lipid assays were performed following a 6-hour fast (starting ∼ZT2), during which mice were housed in clean cages with ad libitum water access. For glucose tolerance tests, fasted mice were injected with glucose (i.p., 1 g/kg). Tail-blood samples were collected at baseline at 15, 30, 45, 90, and 120 minutes after glucose injection. Blood glucose levels were quantified using a hand-held glucometer (Roche, Accu-Check Compact). For triglyceride measurements, plasma was isolated from tail blood and measured (3 μL in technical duplicates) using a commercially available kit (Cayman Chemicals #10010303) as per the manufacturer’s protocols. All plasma samples were stored at -80°C before analysis. Body composition to determine % lean and fat mass was obtained by dual-energy x-ray (DEXA, GE Lunar PIXImus).

Volumetric bone density and bone volume for mice fed SD and HFD were measured at the right femur using a Scanco Medical μCT 50 specimen scanner calibrated to a hydroxyapatite phantom by the Wash U Bone Core. Briefly, samples were fixed in 10% phosphate-buffered formalin and scanned in 70% ethanol. Scanning was performed using a voxel size of 10 mm and an X-ray tube potential of 55 kVp and X-ray intensity of 109 μA. Scanned regions included a 2 mm region of the femur proximal to the epiphyseal plate and a 1 mm region of the femoral mid-diaphysis. Scanned femurs were performed with 10 μm resolution at 70 kV, 57uA, 4W, and an integration time of 700ms. The analysis threshold for cortical and trabecular bone was 0.8 sigma, 1 support, and 260 (lower) and 1000 (upper) permille. Volumes of interest were evaluated using Scanco evaluation software. Representative 3D images were created using Scanco Medical mCT Ray v4.0 software.

### Bone Histomorphometry Analysis

Female mice were injected with 20 mg/kg of calcein (Sigma-Aldrich, St. Louis, MO, USA) 9 days before euthanasia and 15 mg/kg of Alizarin (Sigma-Aldrich) 2 days before euthanasia. Bones were fixed in 4% formalin, dehydrated in 30% sucrose, and embedded in OCT. Standard undecalcified sections (5 mm) were cut using a microtome (Leica CM1950) together with the CryoJane Tape-Transfer System. Mounted sections were imaged with an ECHO REVOLVE R4 using FITC (Calcein) and TexasRed (Alizarin Red) channels by the Wash U Bone Core. A standard sampling site with an area of2.5 mm2 was established in the secondary spongiosa of the distal metaphysis. Prior to histomorphometry analyses, mosaic-tiled images of distal femurs were acquired at ×20 magnification with a Zeiss Axioplan Imager M1 microscope (Carl Zeiss MicroImaging) fitted with a motorized stage. The tiled images were stitched and converted to a single image using the Axiovision software (Carl Zeiss MicroImaGing) prior to blinded analyses being performed using two image-analysis software programs, Bioquant OSTEO software (Nashville, TN, USA) or ImageJ software. The following variables were analyzed: bone volume/total volume (BV/TV), mineral apposition rate (MAR), mineral surface/bone surface (MS/BS), bone formation rate/bone surface (BFR/BS) and osteoclast number/bone surface (Oc/BS). Adjacent sections were stained with Alizarin Red for overview brightfield images.

### TRAP and osmium staining For BMAT Quantification

Femoral samples were demineralized in 10% EDTA for 10–14 days before being embedded in MMA plastic. Sections (5 μm) were cut using the Leica RM2165 and subsequently stained with hematoxylin and eosin (H&E) or stained with Tartrate-resistant acid phosphatase (TRAP) by the Wash U Bone Core. Photoshop software removed the background in non-tissue areas for images of the proximal tibias. H&E and Sirius Red staining were done by the Histology and Microscopy Core Facility at Wash U.

Quantification of BMAT followed published protocol 27. Briefly, femurs and tibiae were decalcified in 14% EDTA, pH 7.4 for 2 weeks, followed by incubation with a PBS solution containing 1% osmium tetroxide (Electron Microscopy Sciences 19170) and 2.5% potassium dichromate (Sigma-Aldrich 24–4520) for 48 hr. After washing for 2 hr with water, osmium-stained bones were embedded in 2% agarose prior to scanning at 10 μm voxel resolution with a Scanco μCT 40 scanner. Regions of interest were contoured and analyzed with a threshold of 400 for BMAT quantification. Specifically, a region of 2 mm right above the growth plate in distal metaphysis was used for the quantification of rBMAT in femurs.

### Biomechanical Strength Testing

Femurs underwent a three-point bend test using the Instron E100 mechanical load frame. A span of 7 mm separated the lower supports to support two ends of the specimen. The testing head was aligned at the midpoint between the supports. Femurs were preloaded to a force of 1 N and then loaded at a rate of 0.1 mm/s. Loading was terminated upon mechanical failure, determined by a drop in force to 0.5 N. Force displacement data was collected every 0.01 s. All tests were performed at room temperature using an electro-mechanical load frame (Instron E1000; Norwood, MA).

### Plasma Collection and Whole Bone Assays

Briefly, 300 μl of whole blood was collected from the submandibular vein from 3-week-old *Esr1*^*fl/fl*^ or *Esr1*^*Nkx2-1Cre*^ female mice into EDTA-treated tubes (Microvette CB 300 K2E) and placed directly on ice. To isolate plasma, whole blood was spun down at 2,000xg for 15 minutes at 4 degrees, and the supernatant was collected. The right and left femurs of 10 to 11-week-old control female and male mice (*Esr1fl/f*) were collected and cleaned of soft tissue. The left femur was fixed in 4% PFA and then transferred into PBS at 4°C for histological assessment for obtaining baseline measurements. The right femur was cultured in a 12-well plate containing1.4 ml of primary culture medium (α-MEM; containing l-glutamine and nucleosides; Mediatech, Herndon, VA, USA), supplemented with 10% FBS (Atlanta Biologicals, Norcross, GA, USA) and 100 U/ml penicillin/streptomycin (Mediatech).

For assessing plasma, the left and right femurs were treated with 15 μl of plasma from *Esr1*^*fl/fl*^ or *Esr1*^*Nkx2-1Cre*^ females, respectively, or with mCCN3 (14 μl of 0.0125 μg/ μl of recombinant mouse (m) NOV/CCN3, Catalog # 1976-NV-050, R&D Systems,) or vehicle (14 μl of 0.9% normal saline), respectively. To assess the degradation of whole bone during culturing, the right tibia or femur was cultured in media with 14 μl of 0.9% normal saline for 5 days (Saline). These were compared to the baseline contralateral tibia or femur, which were immediately chilled and fixed in 4% PFA for analysis (Baseline). Medium changes, including plasma, mCCN3 or vehicle treatments, were performed daily. Femurs were collected after 5 days of culture, fixed in 4% PFA, and then transferred into PBS at 4°C before micro-CT imaging. After micro-CT imaging, femurs were processed for histology as described within the methods.

Femoral samples were cleaned of soft tissue, fixed in 4% PFA, and demineralized in 10% EDTA for 10–14 days before being embedded in paraffin wax. Sections measuring 5 μm were then cut using the Leica RM2165 and subsequently stained with Movat’s Pentachrome staining kit (Abcam Cat# ab245884). For H&E staining analysis, femurs were collected, fixed in 4% formalin, decalcified in Cal-Rite, dehydrated in 30% sucrose, and embedded in OCT. Then, 5 mm standard sections were cut using a microtome (Leica CM1950).

### Brain RNA-Scope and Immunohistochemistry

Cryosections (20 μm) collected from brains fixed in 4% paraformaldehyde were used for both fluorescent ISH and immunostaining. Fluorescent ISH was performed using RNAScope (ACD, Multiplex Fluorescent V2) according to the manufacturer’s protocol using the following probes: *Ccn3*/*Nov* (ACD, #415341-C2) and *Esr1* (ACD, #478201).

Immunohistochemistry was performed using primary antibodies against ERα (EMD Millipore, #C1355 polyclonal rabbit, 1:750 dilution), CCN3 (R&D Systems, #AF1976 polyclonal goat, 1:1000 dilution) and KISS1 (Abcam # ab19028 polyclonal rabbit, 1:200 dilution) diluted in PBS with 0.1% Triton-X100, 5% normal donkey serum, and 5% BSA. For detection, sections were labeled with species-appropriate secondary Alexa Fluor-coupled antibodies (Invitrogen, #A-21447, #A10042, or #A-11055; 1:1000 dilution). Slides were imaged with a Keyence BZ-X800 widefield fluorescence microscope. Confocal images were acquired at the UCSF Nikon Imaging Center using a Nikon CSU-22 with an EMCCD camera and MicroManager v2.0gamma. Images were processed and quantified using ImageJ FIJI v1.52i and the Cell Counter plugin v2. Three representative views of each sample were selected. A complete list of all antibodies used in immunohistochemistry analyses are listed in Extended Data Table 2.

### SiRNA Studies

Mice were secured in a stereotaxic frame (Model 1900, David Kopff Instruments), and 400 nL of *Ccn3* or non-targeting siRNA pools (Dharmacon, E-040684-00-0010 or D-001810-10-05, 0.4 mM) were injected bilaterally to the ARC at the following coordinates: A-P: Bregma -1.58 mm, M-L: Bregma +/-0.25 mm, D-V: skull -5.9 mm. After 12 days post-injection, mice were euthanized, and brain and bone samples were collected and processed as described above. Isolated femurs were then imaged by μCT as described above for bone transplant assays. Due to the high bone phenotype of mutant female mice, thresholding and ROI selection were adjusted between different experiments but kept consistent for individual experiments.

### CCN3, S961, and Naloxone In Vivo Treatments

For recombinant CCN3, 10-week-old control female and 13-weekold control male mice (*Esr1fl/fl)* were injected daily with recombinant mouse NOV/CCN3 at 7.48 μg/kg (R&D Systems Catalog # 1976-NV-050) or vehicle control (0.9% NS). by intraperitoneal (i.p.) administration for 21 days. Seven days before euthanasia, mice were first injected with 20 mg/kg of Calcein (Sigma-Aldrich, St. Louis, MO, USA) and then 15 mg/kg of Alizarin (Sigma-Aldrich, St. Louis, MO, USA) two days before euthanasia. Right femurs were cleaned of soft tissue, fixed in 4% PFA, and then stored in PBS at 4°C prior to imaging. Isolated femurs were imaged by μCT scanning as described above for bone transplant assays. After imaging, femurs were then processed for H&E histology and dynamic histomorphometry as described above.

For S961 treatment, 10-week-old *Esr1*^*fl/fl*^ and *Esr1*^*Nkx2-1Cre*^ female mice were infused with continuous S961 (70 nM per osmotic pump), an insulin peptide receptor antagonist, as adapted from^29^ and obtained from the Novo Nordisk, Compound Sharing Program (Cat# NNC0069-0961) or vehicle solution via an ALZET mini osmotic pumps (Model 1004) which was exchanged once after 4 weeks for a total infusion period of 8 weeks. S961 was reconstituted in 0.9% normal saline. The mini osmotic pumps were implanted into an interscapular subcutaneous pocket under isoflurane anesthesia and exchanged once.

For Naloxone treatments, 10-week-old *Esr1*^*fl/fl*^ and *Esr1*^*Nkx2-1Cre*^ female mice were infused with continuous Naloxone, a non-selective opioid antagonist (0.5 mg/24 hours over 4 weeks, Tocris, Cat# 0599/100) or vehicle solution via an ALZET mini osmotic pump (Model 1004) for 28 days. Naloxone was reconstituted in the vehicle solution containing 0.9% Normal Saline. The mini osmotic pump was implanted into an interscapular subcutaneous pocket under isoflurane anesthesia.

### Hepatic Viral Transductions of mCCN3

Ectopic hepatocyte expression of mCCN3 protein was achieved using AAVdj viral vectors encoding mouse CCN3 under the control of the constitutive cytomegalovirus immediate-early enhancer/ chicken β-actin promoter (CAG) promoter (Vector Biolabs, AAV-265951). Viral vectors were first diluted in sterile saline, and control (*Esr1*^*fl/fl*^) mice were injected retro-orbitally with 100 μL of low (5*10^10^ GC/mL) or high (3*10^11^ GC/mL) titers of AAVdj-CAG-CCN3 or the negative control. To specifically label liver-secreted proteins, mice in the high titer group and their controls were co-injected with AAV-TBG-TurboID (5*10^10^ GC/mL, a gift from Dr. E. Goldberg, UCSF). After 2 weeks, one animal from each group was euthanized, and its liver was checked for expression of CCN3 using an anti-CCN3 antibody as listed in Extended Data Table 2. After 5 weeks, mice were euthanized, and bone, plasma, and liver were collected. Liver samples were divided and processed separately for analysis of *Ccn3* mRNA expression by qPCR or CCN3 protein expression by immunohistochemistry. Total liver RNA was isolated by phenol-chloroform extraction and purified using the RNeasy Mini Kit (Qiagen, #74104). qPCR was performed as described below. For immunohistochemistry, liver samples were drop-fixed in 4% PFA, cryosectioned (10 μm), and antibody stained as described above. Isolated femurs were cleaned and then imaged as described above for bone transplant assays.

### RNA isolation, qPCR and bulk RNA-Seq

Microdissected ARC or medial basal hypothalamic tissue was obtained from control and mutant female mice (1-27 weeks of age) using the optic chiasm as a reference point, and a 2 mm block of tissue containing the hypothalamus was isolated with a matrix slicer. For ARC, total RNA was purified using the RNA Mini Kit (Invitrogen, Waltham MA). For quantitative polymerase chain reaction (qPCR), cDNA was synthesized using the Applied Biosystems High-Capacity cDNA Reverse Transcription Kit. expression analysis was performed using SYBR Green. Values were normalized to either *36b4, mCyclo*, or *Gapdh*. Sequences for primer pairs can be found in Extended Data Table 1.

For bulk-RNA-Seq analyses, barcoded sequencing libraries were prepared from RNA samples after a quality check (QC), and sequencing was performed via a paired-end 150 bp strategy where single-end 50 bp reads were sequenced from the multiplexed libraries; these steps were carried out by Novogene (Davis, CA). For all tissue samples, sequencing-generated reads were aligned to the mouse transcriptome (mm10) using Kallisto in gene mode^47^. Differential gene expression was evaluated using the likelihood-ratio test by Sleuth (qval <0.05)^48^. All heatmaps were generated with the top 50 female/male-biased genes obtained from 27-week-old mice and were generated in R^49^.

### Plate-based SmartSeq2 Single-Cell RNA-sequencing of ocSSCs

Single ocSSCs from 4-week-old *Esr1*^*fl/fl*^ or *Esr1*^*Nkx2-1Cre*^ long bones were isolated via FACS using processing and flow cytometry protocol as described above. Single-cell suspension of four mice per group were pooled, and individual cells were captured in separate wells of a 96-well plate containing 4 μl lysis buffer (1 U/μl RNase inhibitor [Clontech, Cat#: 2313B]), 0.1% Triton (Thermo Fisher Scientific, Cat#: 85111), 2.5 mM dNTP (Invitrogen, Cat#: 10297–018), 2.5 μM oligo dT30VN (IDT, custom: 5′–*AAGCAGTGGTATCAAC-GCAGAGTACT30VN*-3′), and 1:600,000 External RNA Controls Consortium ExFold RNA Spike-In Mix 2 (ERCC; Invitrogen, Cat#: 4456739) in nuclease-free water (Thermo Fisher Scientific, Cat#: 10977023) according to a modified SmartSeq2 protocol^46^,^50^. Two 96-well plates per phenotype with a single ocSSCs per well were sorted and processed. Plates were spun down and kept at −80°C until complementary DNA (cDNA) synthesis, which was conducted using oligo-dT-primed reverse transcription with SMARTScribe reverse transcriptase (Clontech, Cat#: 639538) and a locked nucleic acid containing template-switching oligonucleotide (TSO; Exiqon, custom: 5′-*AAGCAGTGGTATCAACGCAGAGTACATr-GrG*+G-3′). PCR amplification was conducted using KAPA HiFi HotStart ReadyMix (Kapa Biosystems, Cat#: KK2602) with In Situ Polymerase Chain Reaction (ISPCR) primers (IDT, custom: 5′-*AAGCAGTGGTATCA-ACGCAGAGT*-3′). Amplified cDNA was then purified with Agencourt AMPure XP beads (Beckman Coulter, Cat#: A63882). After quantification, cDNA from each well was normalized to the desired concentration range (0.05–0.16 ng/μl) by dilution and consolidated into a 384-well plate. Subsequently, this new plate was used for library preparation (Nextera XT kit; Illumina, Cat#: FC-131–1096) using a semi-automated pipeline. The barcoded libraries of each well were pooled, cleaned-up, and size-selected using two rounds (0.35x and 0.75x) of Agencourt AMPure XP beads (Beckman Coulter), as recommended by the Nextera XT protocol (Illumina). A high-sensitivity fragment analyzer run was used to assess fragment distribution and concentrations. Pooled libraries were sequenced on NovaSeq6000 (Illumina) to obtain 1–2 million 2 × 151 base-pair paired-end reads per cell.

### Single-cell RNAseq data processing

Sequenced data were demultiplexed using bcl2fastq2 2.18 (Illumina). Raw reads were further processed using a skewer for 3′ quality trimming, 3′ adaptor trimming, and removal of degenerate reads. Trimmed reads were then mapped to the mouse genome vM20 using STAR 2.4, and counts for gene and transcript reads were calculated using RSEM 1.2.21. Data were explored, and plots were generated using Scanpy v1.9. To select high-quality cells only, we excluded cells with fewer than 450 genes and genes detected in less than three cells were excluded. Cells with a mitochondrial gene content higher than 5%, ERCC content higher than 30%, and ribosomal gene content higher than 5% were excluded as well. Scrublet was then used to detect and remove residual duplicates. A total of 264 high-quality cells (122 control and 142 mutant mouse cells) were included in the final analysis. Raw counts per million (CPM) values were mean- and log-normalized, and then data were scaled to a maximum value of 10. Combat batch correction was applied to account for potential biases through minor differences in cell processing. Principal component (PC) ‘elbow’ heuristics were used to determine the number of PCs for clustering analysis with UMAP and Leiden Algorithm (leidenalg). Differential gene expression between *Esr1*^*fl/fl*^ (wild type) and *Esr1*^*Nkx2-1Cre*^ (mutant), as well as Leiden clusters, was calculated by the Wilcoxon-Rank-Sum test. Cell cycle status was assessed using the ‘score_genes_cell_ cycle’ function with the updated gene list provided by Nestorowa et al.^51^. EnrichR was used to explore enrichment for pathways and ontologies of differentially expressed genes between wild-type and mutant groups^52^.

### Statistics

Statistical tests, excluding RNA-Seq analyses, were performed using Prism 10 (GraphPad). A description of the test and results are provided in Extended Data Table 3. Multiple comparisons correction for One-way, Two-way, and repeated measures (RM) ANOVA were performed using the Šidák post hoc test. For all panels in the Main and Extended Data Figures, N = biological sample sizes used and n = technical replicates in cell culture assays. Unless otherwise noted, data are presented as mean ± SEM or box plots in which whiskers represent minimum and maximum values, edges of the box are 25th and 75th percentiles, and the center line indicates the mean. Sample sizes are based on previous work from our labs; however, no specific statistical calculation was performed to determine sample size. For AAVdj-CCN3 and siRNA injections, mice of identical genotypes were drawn at random from littermate pools to receive functional or control virus injections. Experimenters were blinded to the type of AAV received/genotype of the mice under study for subsequent μCT and dynamic histomorphometry analyses. All raw data and processed data files for the bulk-RNA and sc-RNA Sequencing are publicly available at GEO under sample accession number GSE241478.

## ACKNOWLEDGMENTS

We thank Ms. J. Argiris for initial expert assistance with animal husbandry and genotyping and the UCSF Nikon Imaging Core for the use of their confocal imaging facilities. We thank all members of our groups for helpful suggestions and critical comments and for strategic contributions in the early stages of this project, including D. Julius, as well as. Raghav Kalia for his structural insights into CCN3 and its potential mechanism of action.

This work was supported by NIH Transition Awards including an NIDDK Emerging Physician Award R01DK121657-S1 (M.B.), an NIA-1K01AG065916 (C.B.H.), the NIGMS K12 IRACDA 5K12GM081266 (R.R.), an NIDDK K99DK129763 (J.N.), and an NIA K99/R00 AG066963 (T.A.). Grants from the USA NIH include R01 support NDDK R01DK132073 (E.L.S.), R01AG067740 (S.V.), NIA R01AG070647 (N.E.L.) and R01AG062331, R01DK121657 (H.A.I.). Non-NIH Support came from a Stanford Pilot Award (H.A.I, C.F.C.) and a Senior Scholar Award GCRLE0320 (H.A.I.). Core services were made possible by funding from NIH NIAMS Support for Cores P30AR075055 (at UCSF, School of Medicine) and P30AR075055 (at Washington University).

## DATA AVAILABILITY STATEMENT

All data generated or analyzed during this study will be included in the published article (and its supplementary information files).

## AUTHOR CONTRIBUTIONS

M.E.B, W.C.K., C.B.H. T.H.A., and H.A.I. conceived and designed the experiments, interpreted the results, and wrote the paper. M.B. designed and conducted ex-vivo bone assays and in vi treatments with drugs and HFD. W.C.K. performed all brain histology, stereotaxic surgery, and viral CCN3 injections. C.B.H. conducted parabiosis studies with S.V., as well as whole bone transplants, ex-vivo whole bone, and ocSSC culturing. K.C. conducted bone histology and histomorphometry analyses on bone following viral CCN3 transductions. T.H.A. quantified all bone parameters except for mice treated with SD and HFD and performed scSeq RNA Seq analyses and ocSSCs cell assays. J.N. performed analyses on bulk RNA-Seq data sets. R.R. performed stereotaxic surgery of isolated ocSSCs into the brain. F.C.N. and Y.W. provided excellent technical support on multiple aspects of this project. X.Z. and E.L.S. determined BMAT levels in mouse models. N.E.L. and C.F.C. provided guidance on the project, including bone expertise and bone stem cell niches, and edited the manuscript.

## EXTENDED DATA FIGURES AND TABLES

Data will be provided upon request.

